# Intrinsic spine dynamics are critical for recurrent network learning in models with and without autism spectrum disorder

**DOI:** 10.1101/525980

**Authors:** James Humble, Kazuhiro Hiratsuka, Haruo Kasai, Taro Toyoizum

## Abstract

It is often assumed that Hebbian synaptic plasticity forms a cell assembly, a mutually interacting group of neurons that encodes memory. However, in recurrently connected networks with pure Hebbian plasticity, cell assemblies typically diverge or fade under ongoing changes of synaptic strength. Previously assumed mechanisms that stabilize cell assemblies do not robustly reproduce the experimentally reported unimodal and long-tailed distribution of synaptic strengths. Here, we show that augmenting Hebbian plasticity with experimentally observed intrinsic spine dynamics can stabilize cell assemblies and reproduce the distribution of synaptic strengths. Moreover, we posit that strong intrinsic spine dynamics impair learning performance. Our theory explains how excessively strong spine dynamics, experimentally observed in several animal models of autism spectrum disorder, impair learning associations in the brain.

## Introduction

The operation of a neural circuit is shaped by the strength of synapses that mediate signal transduction between neurons. Activity-dependent modification of synaptic strength, termed synaptic plasticity, is considered to be an underlying mechanism of learning and memory (Malenka and Bear 2004; Mongillo et al. 2017). A major form of synaptic plasticity is Hebbian plasticity (Hebb 1949). While there are multiple molecular mechanisms (Malinow and Malenka 2002; Nicoll et al. 2006; Matsuzaki et al. 2004) underlying Hebbian plasticity and experimental protocols (Neves et al. 2008), it is commonly induced by coactivation of pre- and postsynaptic neurons within a particular time window. One prominent biological mechanism for Hebbian plasticity is activity-dependent spine volume change. Spine volume is known to be tightly correlated with synaptic strength (Matsuzaki et al. 2001; Smith et al. 2003; Noguchi et al. 2005; Béïque et al. 2006; Asrican et al. 2007; Holbro et al. 2009; Zito et al. 2009), and both long-term potentiation (LTP) and long-term depression (LTD) involve spine change (Lang et al. 2004; Matsuzaki et al. 2004; Otmakhov et al. 2004; Zhou et al. 2004; Kopec et al. 2006; Hayama et al. 2013).

It was previously proposed (S. I. Amari 1977; Hopfield 1982; Hebb 1949) that a memory can be represented by coherent activity in a cell assembly, i.e., a group of cells mutually exciting each other, and the memory can be stored in synaptic strengths between these neurons by Hebbian plasticity. Consistently, recent experiments have shown that the activation of a coherently active group of cells is necessary and sufficient for the expression of learned behavior (Liu et al. 2012; Nabavi et al. 2014). However, how a neural circuit maintains cell assemblies stably is not well understood. In some models, cell assemblies are stable because synaptic strength is modified only during learning, and then fixed (Hopfield 1982; Vogels et al. 2011; S. Amari 1977). However, these models neglect changes in synaptic strength after learning and thus do not address the maintenance of an acquired memory.

Several studies modeled ongoing Hebbian plasticity during spontaneous activity and found that Hebbian plasticity alone is likely not sufficient to maintain a cell assembly. In additive Hebbian plasticity models (Song et al. 2000; Gütig et al. 2003; Gerstner et al. 1996), in which the dependencies of the LTP and LTD amplitudes on synaptic strength are the same, memory tends to become unstable due to a positive feedback during spontaneous activity (Litwin-Kumar and Doiron 2014; Zenke et al. 2015; Fiete et al. 2010), namely, neurons that fire together are wired together, and then fire together more often. This kind of a positive feedback process typically fuses assemblies, and expands the largest existing cell assembly. Some forms of stabilizing mechanisms, such as inhibitory plasticity (Litwin-Kumar and Doiron 2014), homeostatic plasticity (Zenke et al. 2013), or heterosynaptic plasticity (Zenke et al. 2015) have been suggested to stabilize memory (Keck et al. 2017). However, even with these stabilizing mechanisms, the resulting distribution of synaptic strengths often becomes dissimilar to what has been experimentally observed (Toyoizumi et al. 2007). For example, while the models with positive feedback often produce a synaptic strength distribution that is bimodal, experiments have reported a unimodal and long-tailed distribution of synaptic strengths (Song et al. 2005; Cossell et al. 2015) and corresponding spine volumes (Yasumatsu et al. 2008; Loewenstein et al. 2011).

An alternative proposal is multiplicative Hebbian plasticity (van Rossum et al. 2000; Morrison et al. 2007; Gütig et al. 2003), in which the LTP amplitude is less prominent than the LTD amplitude for large synapses, in agreement with experimental observations (Bi and Poo 1998; Tanaka et al. 2008; Hayama et al. 2013). This multiplicative form of Hebbian plasticity can avoid the above instability problem, and under spontaneous activity of neurons, synaptic strengths converge to a prefixed set point where LTP and LTD effects balance each other, regardless of the initial synaptic strengths (Morrison et al. 2007). This means that memories must degrade in the presence of spontaneous neural activity.

Hence, in all the models described above, it is nontrivial to stably maintain cell assemblies and reproduce the experimentally observed distribution of synaptic strengths (or of spine volumes), which has a thick tail and a peak at a rather weak strength (Song et al. 2005; Yasumatsu et al. 2008; Loewenstein et al. 2011; Cossell et al. 2015). Interestingly, a similar distribution of spine volumes is robustly observed even in animal models of mental disorders (Pathania et al. 2014) and with an LTD deficiency in calcineurin KO animals (Okazaki et al. 2018). In contrast, in the above mathematical models, the distribution of synaptic strengths is fragile and strongly depends on the balance of LTP and LTD that is set by model parameters and input to neurons. Thus, conventional models have no mechanism to restore the distribution of synaptic strengths.

Despite the common assumption that only synaptic plasticity changes spines, they also dynamically change in the absence of neural activity. Recent studies showed that spine turnover, i.e., generation and elimination of spines, continues even under the blockade of neural activity and calcium signaling *in vivo (Kim and Nabekura 2011; Nagaoka et al. 2016)*. Further, spine volumes constitutively fluctuate in the absence of neural activity, calcium signaling, and activity-dependent plasticity *in vitro* (Yasumatsu et al. 2008). These intrinsic spine dynamics are characterized by a zero drift coefficient and a diffusion coefficient proportional to the square of spine volume, *v*^2^. Interestingly, this volume-dependent diffusion reproduces the experimentally observed equilibrium distribution of spine volumes with a power-law tail of exponent *v*^−2^ (Yasumatsu et al. 2008; Ishii et al. 2018). This observation poses an important question: How do intrinsic spine dynamics affect the maintenance of cell assemblies?

We address this question by simulating a mathematical model of a recurrently connected neural network that implements both multiplicative spike-timing dependent plasticity (STDP) (van Rossum et al. 2000; Morrison et al. 2007) and experimentally observed intrinsic spine dynamics (Yasumatsu et al. 2008). We also study how spine turnover and the distribution of spine volumes are affected by these two processes. Despite a possible perception of intrinsic spine dynamics as *noise*, we show that they can help to maintain cell assemblies by preventing unnecessary spines from growing and sustaining the physiological spine volume distribution.

Based on the model analysis, we hypothesize that intrinsic spine dynamics that are stronger than in wild type (WT) conditions can explain the abnormally high spine turnover rate observed in animal models of autism spectrum disorder (ASD) (Isshiki et al. 2014; Pan et al. 2010). By fitting model parameters to one ASD mouse model, *fmr1*KO (a model of fragile X syndrome) (Pfeiffer and Huber 2009), we show how excessively strong intrinsic spine dynamics may cause learning deficits in ASD animals (Silverman et al. 2010; Padmashri et al. 2013).

## Results

We study the dynamics of spine volumes using a model of a cortical circuit. We stimulate recurrently connected spiking neurons (Fig. 1; also see Methods) to explore if the network stores memories as cell assemblies. The cortical network consists of 1000 excitatory and 200 inhibitory leaky-integrate-and-fire spiking neurons (Tuckwell 1988), where the excitatory and inhibitory neurons are randomly connected by a 10% connection probability (Fig. 1A). For simplicity, the excitatory neurons are embedded in a one-dimensional feature space that describes, for example, orientation selectivity in V1. We assume that synapses can be formed between a pair of excitatory neurons that have potential connectivity (Markram et al. 1997). Potential connectivity (either 0 or 1) from a neuron to another is randomly generated and set at the beginning of a simulation. Potential connectivity peaks at 10.4% (see Fig. 7 for a systematic exploration of this peak value) for neurons with similar selectivity and falls off with their tuning-distance (Fig. 1B_1_). If two neurons have potential connectivity, the number of synaptic contact points is drawn randomly from a truncated Poisson distribution (Fig. 1B_2_) (Hardingham et al. 2010). In the absence of elevated external input, excitatory neurons in this network exhibit a background firing rate of about 0.1 Hz (Fig. 1B_3_). Cortical excitatory neurons are generally adaptive and cannot continuously fire at their maximum firing rate. Hence, we model an adaptation current (Wang et al. 2003) that slowly builds up with the postsynaptic spiking activity and hyperpolarizes the neuron with its characteristic time constant of about 5 s (Fig. 1B_4_; see Methods). Finally, only the recurrent excitatory to excitatory synapses are subject to activity-dependent plasticity.

**Figure 1:**
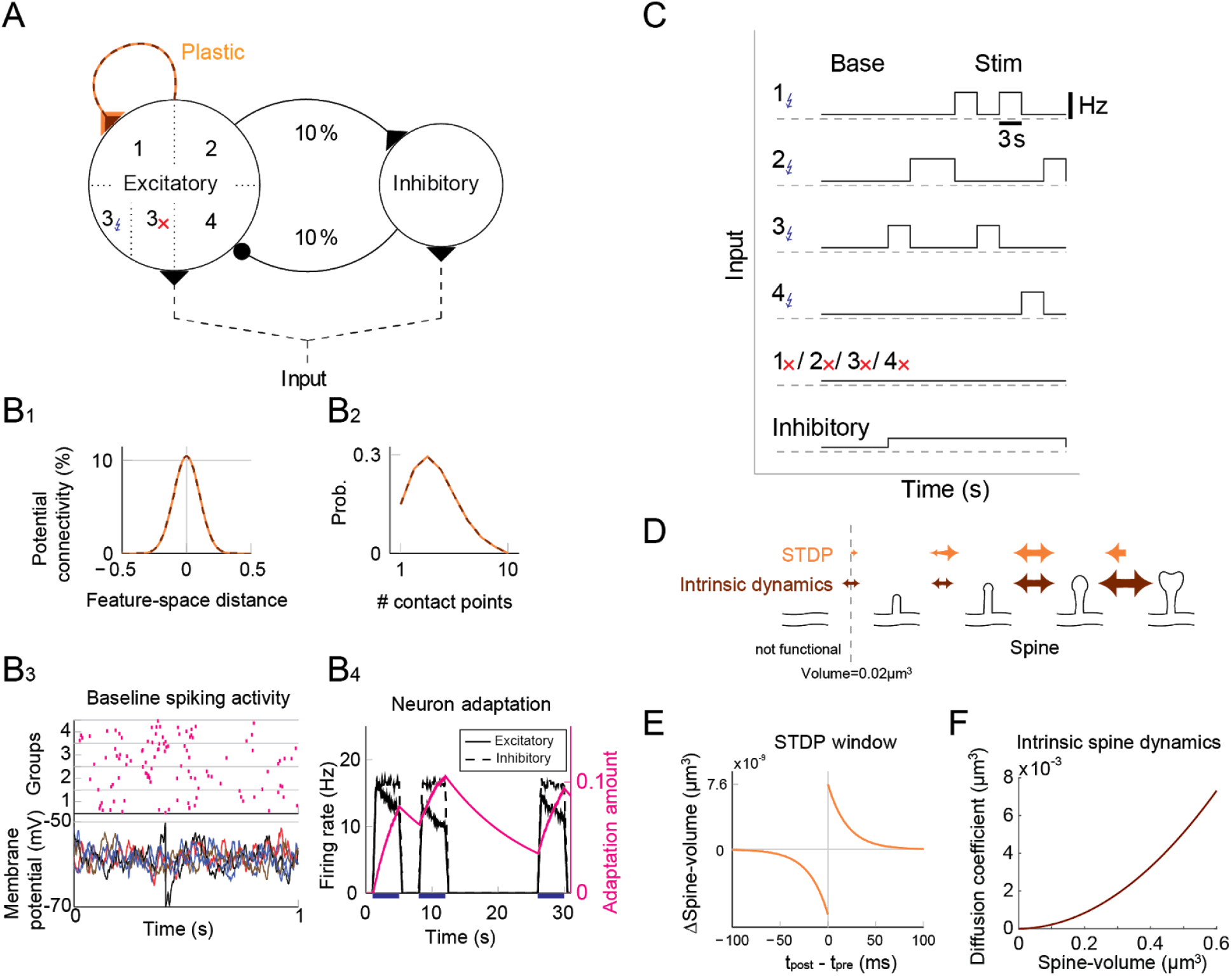
Recurrent network model with STDP and intrinsic spine dynamics. (A) A model of cortical circuitry. Excitatory and inhibitory neurons are modeled as leaky-integrate-and-fire units, which are sparsely connected. We assume that only the recurrent excitatory synapses are plastic. Excitatory neurons are aligned in a one-dimensional feature space, which is divided into 4 neighborhood quarters. 40% of randomly chosen excitatory neurons in each group, and all inhibitory neurons, receive additional external input during stimulation. The externally stimulated excitatory neurons in each quarter are defined as a stimulated group. (B_1_) Potential connectivity peaks at around 10% and decays with the tuning-distance between two excitatory neurons in the feature space. Synapses can grow if two excitatory neurons are potentially connected. (B_2_) If two excitatory neurons have potential connectivity, the number of contact points is randomly drawn from a truncated Poisson distribution in the range of 1 to 10. Each contact point can accommodate one spine. (B_3_) Spiking activity and membrane potential dynamics of a sample set of neurons at baseline. (B_4_) Excitatory neurons have an adaptation current, which builds up with firing activity and suppresses firing rate. (C) During a learning period, one of the stimulated groups is randomly chosen with probability ¼ and receives elevated external input for 3 s. All inhibitory neurons receive external stimulation throughout the entire learning period. (D) Spine volumes are changed by the combination of STDP and intrinsic spine dynamics (except in Fig. 2, where only STDP is considered). Arrows indicate possible changes in spine volume and the size of the arrow represents the possible maximum change in spine volume. A threshold at 0.02 μm^3^ separates spines and non-spines, and STDP only affects spines. (E) The multiplicative STDP rule used for changing spine volume. The LTD amplitude is proportional to spine volume. (F) The diffusion coefficient characterizing intrinsic spine dynamics, which is proportional to the square of spine volume.

To model activity-dependent plasticity, we assume that spine volume is proportional to synaptic strength (but we define a tiny protrusion of volume <0.02 μm^3^ as a “non-spine”) because the correlation between the synaptic strength and spine volume has been experimentally demonstrated (Matsuzaki et al. 2004; Harvey and Svoboda 2007; Bosch et al. 2014). The spine volume of an excitatory to excitatory synapse is modeled by multiplicative STDP (Fig. 1E) (van Rossum et al. 2000), and thus the LTP amplitude is independent of synaptic strength, while the LTD amplitude is proportional to synaptic strength. Therefore, with an increase in synaptic strength, the LTD amplitude increases at a steeper rate than the LTP amplitude, and this is consistent with experimental observations (Tanaka et al. 2008; Hayama et al. 2013; Bi and Poo 1998). We let the network acquire cell assemblies by providing additional external input to subsets of neurons. We divide the feature space of the excitatory network into 4 equally sized neighboring quarters (Fig. 1A), and randomly select 40% of the neurons in each quarter as a stimulated group. We randomly select one of these four groups at a time during the learning period and stimulate it with an elevated rate of Poisson spikes for 3 s (Fig. 1C; see Methods). For simplicity, all inhibitory neurons are stimulated throughout the learning period. The spine volume, *v*, of each spine is initially drawn randomly from a fixed distribution, proportional to (*v* + 0.05 μm^3^)^−2^. This initial distribution approximates an experimentally observed spine volume distribution (see Methods). As we will see below, changing synaptic strengths by multiplicative STDP alone fails to sustain cell assemblies in the presence of spontaneous activity.

Firstly, we consider the case where spine fluctuations are absent and spine volumes are only changed by multiplicative STDP. Figure 2 depicts the behavior of our network during and after the learning period. The learning period (Fig. 2A) is terminated when the mean intra-group spine volume of at least one stimulated group reaches ≥0.49 μm^3^. During the learning period, four groups of neurons were randomly stimulated one at a time (Fig. 2B, Left) with increased input, and after the learning period, only the 3rd-quarter’s neurons stayed active (Fig. 2B, Right). Figure 2C plots the firing rates of all neurons in the 3rd-quarter during the entire simulation period. This shows that the number of active neurons monotonically increased both during the learning period and during the maintenance period, indicating an unstable learning outcome. Specifically, the cell assembly initially formed among the group 3 neurons and spread to neighboring neurons that were not externally stimulated. This spreading of the cell assembly provided extra recurrent input from newly recruited neurons to other neighboring neurons. Figure 2D summarizes the population averaged firing rates of the four stimulated groups. All groups show elevated firing rates during the learning period due to external stimulations. After the learning period, physiological neural activity was maintained only for a day. After that, the mean firing rate of one group (i.e., group 3) exploded, and that of the other groups declined down to near zero values. The spine volume distribution at the end of the simulation included a non-physiological secondary peak at 0.5 μm^3^ (Fig. 2E). A small but non-negligible number of non-stimulated synapses also formed a peak at 0.5 μm^3^. These synapses contribute to the extremely high firing rate of the group 3 neurons and the recruitment of non-stimulated neurons toward the end of the simulation. During the learning period, the mean volume of the spines connecting neurons within each stimulated group increases due to LTP (Fig. 2F). Note that LTP is dominant over LTD for small spines because we assume that the LTP amplitude is fixed but the LTD amplitude is proportional to spine volume. After a few days of learning, mean spine volumes plateaued at around 0.5 μm^3^, where LTP and LTD effects roughly balance. The mean spine volume of other spines (non-stimulated) exhibited a slow but steady increase (Fig. 2F), reflecting the formation of the secondary peak of non-stimulated spines seen in the spine volume distribution (Fig. 2E). The volumes of individual spines (Fig. 2G) are homogeneous within each group and their development mirrors the corresponding mean spine volume (Fig. 2F).

**Figure 2.**
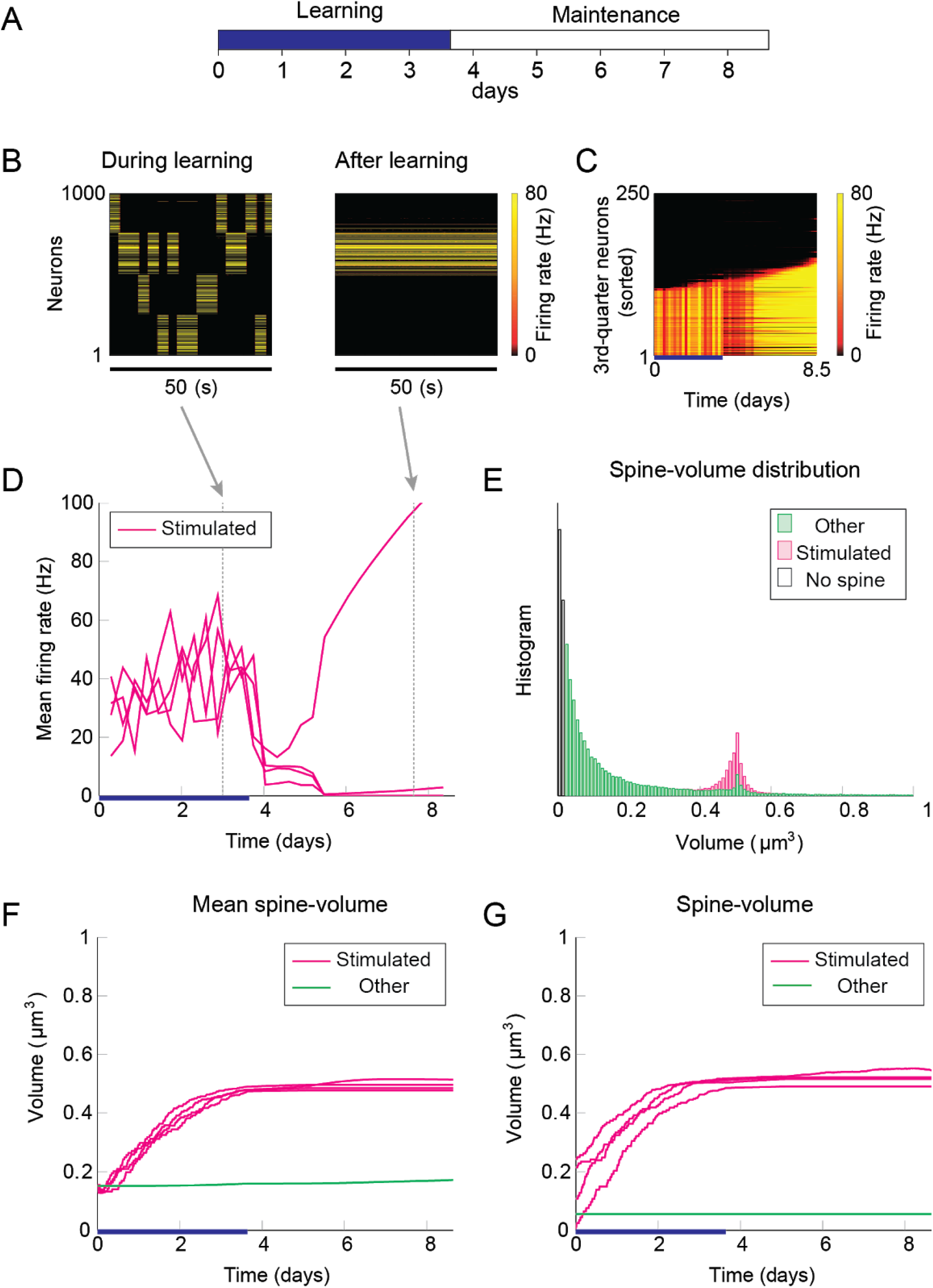
Network behavior in the absence of intrinsic spine dynamics. (A) Experimental protocol. External stimulus is provided during the learning period (blue bar) and the memory retention is studied in the maintenance period. The learning period finishes when one cell assembly becomes strong enough to sustain it’s activity. (B) Typical neural activity during (*Left*) and after (*Right*) the learning period. The panels show the firing rates of 1000 excitatory neurons in a 50 s time window. During the learning, neural activity is driven by external stimulus, which randomly activates one group at a time. After learning, one group of neurons is strongly active spontaneously. (C) Firing rates of the 3rd-quarter neurons in the entire simulation period. Neurons are sorted by the first time their firing rates exceed 15 Hz. The number of active neurons expands both during the learning and retention periods beyond the initial 40 % that are stimulated. (D) Mean firing rates of the four stimulated groups. (E) Spine volume distribution at the end of the simulation for intra-stimulated-group spines (Stimulated: pink) and other spines (Other: green). (F) Mean spine volume of intra-stimulated-group spines (pink) and other spines (green). (G) Individual volumes of a single intra-stimulated-group spine from each stimulated group and a single non-stimulated spine. The learning period is represented by a blue bar.

The spread of activity to non-stimulated neurons is slow in this simulation because of the lateral inhibition that tends to shut down spikes in non-stimulated neurons. However, the activity will eventually spread as long as they fire occasionally. Hence, as expected from previous studies (Morrison et al. 2007), a recurrently connected network with multiplicative STDP has difficulty preventing the expansion of a dominant cell assembly during spontaneous activity. Notably, the resulting spine volume distribution of this model is dissimilar to experimentally observed unimodal distributions.

Next, we add intrinsic spine dynamics previously observed in hippocampal slices (Yasumatsu et al. 2008). Spine volumes fluctuate every day, and the amplitude of these fluctuations is spine volume dependent (Fig. 1F). Importantly, these intrinsic spine dynamics were largely intact in the absence of neural activity and synaptic plasticity, i.e., under the pharmacological blockade of sodium channels, NMDA, and voltage-dependent calcium channels in cells. In the absence of neural activity and plasticity, the amplitude of spine volume fluctuations is roughly proportional to spine volume, ~ α*v* + β, where *v* is spine volume with parameters α = 0.2 day^−½^ and β = 0.01 μm^3^ day^−½^ (Yasumatsu et al. 2008). In other words, the effect of intrinsic spine dynamics is summarized by the volume-dependent diffusion coefficient *D*(*v*) = (α*v* + β)^2^/2 with zero drift coefficient (Yasumatsu et al. 2008) (see also Methods). Therefore, the Fokker-Planck equation (Risken 1989) for describing the evolution of the spine volume distribution *P*(*v*) is 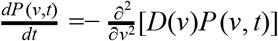. It indicates that the equilibrium is reached when the diffusion intensity *D*(*v*)*P_eq_*(*v*) becomes volume-independent. This gives the equilibrium spine volume distribution *P_eq_*(*v*) ∝ 1/*D*(*v*) ~ *v*^−2^, which has a power-law tail. Note that there are two mathematical conventions for interpreting the above equation (Gardiner 1985), which lead to different semantic meanings of *fluctuation*. Here, we take the Itô interpretation, in which the intrinsic spine dynamics are interpreted as spine volume *fluctuations* (but see Methods for an alternative interpretation). We regard that spines smaller than 0.02 μm^3^ are non-spines and do not exhibit multiplicative STDP, whereas intrinsic spine dynamics are still present even for these small protrusions (*c.f*. Fig. 1D). This assumption is consistent with the experimental observations that the baseline spine turnover is largely activity-independent (Kim and Nabekura 2011; Nagaoka et al. 2016).

We used the same stimulation protocol as in Fig. 2 to study cell assembly learning when both multiplicative STDP and intrinsic spine dynamics are involved (Fig. 3A-H). Cell assembly learning was similar at the beginning of the simulation to the case without intrinsic spine dynamics: Firing rates increased (Fig. 3D) and the intra-group spines enlarged (Fig. 3F). In contrast to the previous case, a physiological neural activity level was maintained throughout (Fig. 3D), and none of the cell assemblies aggressively spread to neighboring non-stimulated spines (Fig. 3C) during the maintenance period. The activity-dependent formation of cell assemblies is evident from the coherent reactivation of stimulated groups (Fig. 3B) during the maintenance period at much lower spontaneous firing rates than during the learning period. These memory patterns can be successfully maintained even if neural activity is blocked for a whole day (e.g., by tetrodotoxin; Fig. S1). Notably, while individual spines’ volumes fluctuated throughout the simulation (Fig. 3G), the mean volume of the intra-group spines was stably maintained (Fig. 3F). In contrast to the previous case without intrinsic spine dynamics (c.f. Fig. 2E), the spine volume distribution remained unimodal after learning, with no secondary peak around 0.5 μm^3^ (Fig. 3E). This is because large spine fluctuations smear large spines along the tail. Despite this smearing, the memory patterns were still stored by enlarged spines located around the fat tail of the distribution. Finally, spine generation was increased during the learning period, as often seen experimentally (Yang et al. 2009; Hofer et al. 2009) (Fig. 3H). The contribution of each model mechanism assumption is further explored in Fig. S2.

**Figure 3.**
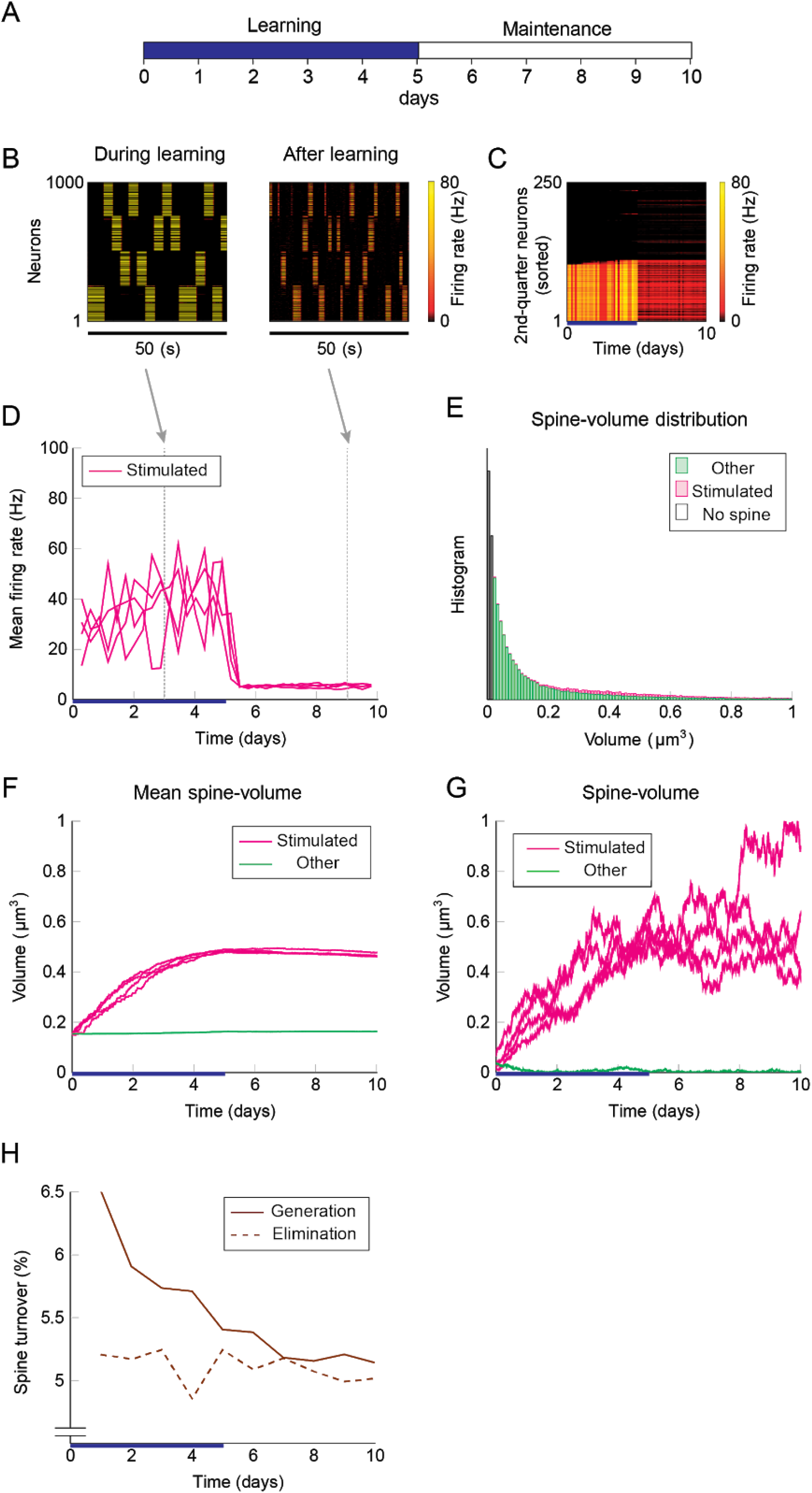
Network behavior in the presence of intrinsic spine dynamics. (A-G) Conventions are as in Fig. 2. (H) Spine generation and elimination.

To elucidate how intrinsic spine dynamics stabilize our network learning and enable the storing of memories, we examined the effects of STDP and intrinsic spine dynamics separately (Fig. 4). We initialized spine volumes randomly as described above except for the intra-group spines of one group, whose volumes were all set identically and changed systematically. While we simulated the spontaneous activity of neurons, we monitored how multiplicative STDP and spine fluctuations changed the mean intra-group spine volume. Multiplicative STDP leads to an overall potentiation of the intra-group spines (Fig. 4; orange line) if they are small (roughly <0.57 μm^3^) and a depression of the spines if they are large (roughly >0.57 μm^3^). This transition is observed for large spines because the LTD amplitude, which is proportional to spine volume, dominates over the amplitude of LTP, which is independent of spine volume. LTP dramatically increases for spines greater than 0.30 μm^3^ because strong intra-group connections serve as positive feedback on coincident firing between the neurons within the group, increasing the frequency of both LTP and LTD events. Therefore, STDP on its own, leads to one fixed point of mean spine volume at a non-physiologically high value at around 0.57 μm^3^. This fixed point is controlled by the parameter setting the relative amplitude of LTD (Morrison et al. 2007). Intrinsic spine dynamics on the other hand normalize the spine volume distribution, restoring the distribution toward the equilibrium distribution *P_eq_*(*v*) ∝ 1/*D*(*v*) with a mean spine volume of roughly 0.15 μm^3^ (Fig. 4; brown line).

**Figure 4.**
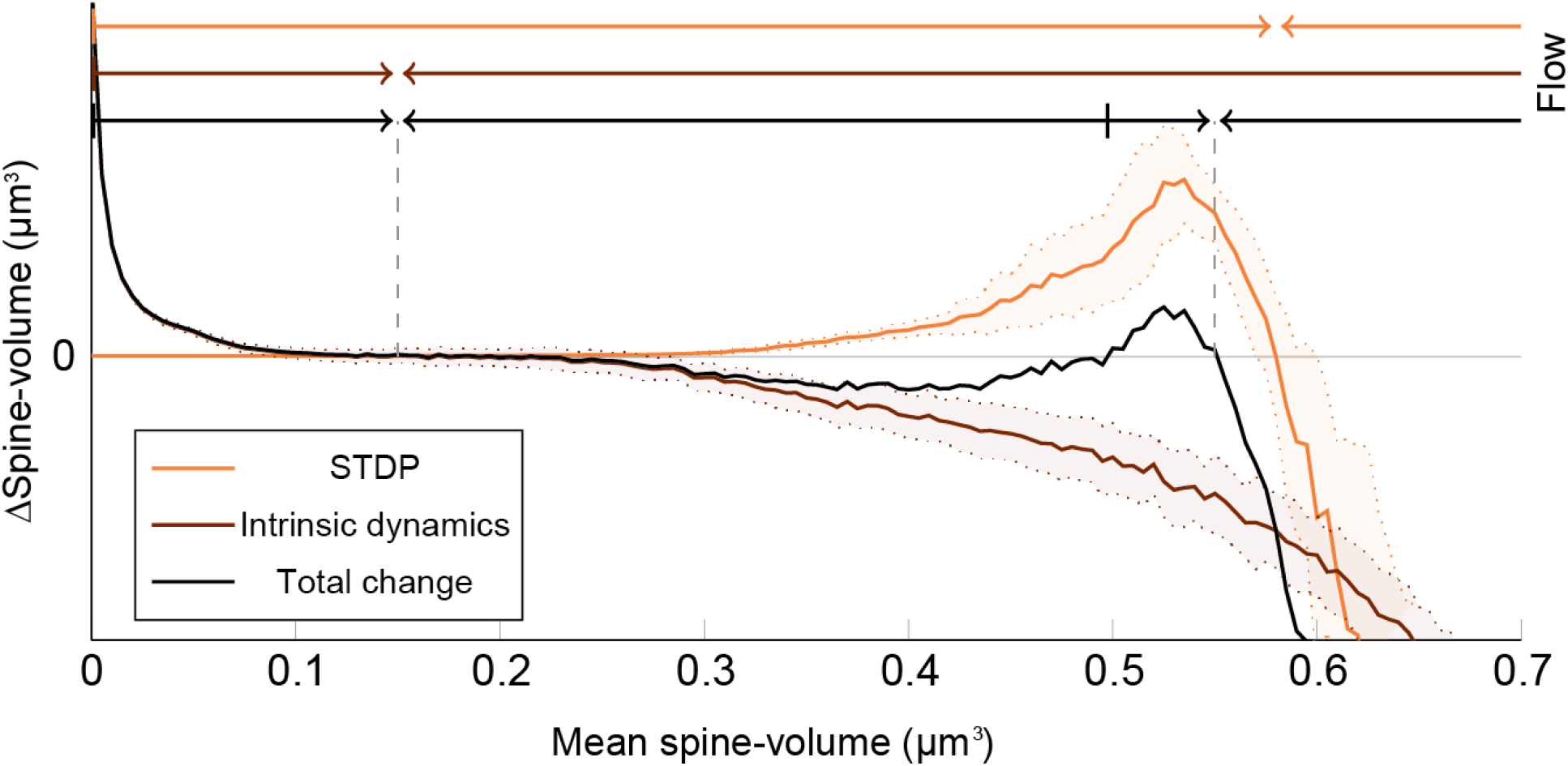
Decomposition of spine volume change by STDP and intrinsic spine dynamics. We separately measured changes in mean spine volume induced by either STDP (orange line) or intrinsic spine dynamics (brown line), by systematically initializing all intra-group spine volumes of one group to a fixed value, and measuring any subsequent changes. The net change is separately plotted (black line). Arrows of corresponding color mark the flow of mean spine volume change due to each mechanism, or the combination of both, at the top. STDP and intrinsic spine dynamics change the mean spine volume toward 0.57 μm^3^ and 0.15 μm^3^, respectively. When the two mechanisms are combined, the net dynamics have bistability: There are two stable fixed points at 0.55 μm^3^ and 0.15 μm^3^, and a separation point at roughly 0.50 μm^3^, which divides the two basins of attraction. The shaded interval indicates the standard deviation of the intra-group spines’ change in the initialized group.

When the contributions from STDP and intrinsic spine dynamics are added together with an appropriate balance (Fig. 4; black line), the combination permits a bi-stability in the mean spine volume of a population of spines. In this case, changes in the mean spine volume are dominated by the intrinsic spine dynamics when small (roughly <0.30 μm^3^), so that the spine volumes fluctuate around 0.15 μm^3^. Conversely, changes in the mean spine volume are significantly affected by STDP when large (roughly >0.30 μm^3^), creating a larger-volume fixed point at approximately <0.55 μm^3^. In between these stable fixed points, there is a separation point (unstable fixed point) at around 0.50 μm^3^, which prevents the mean spine volume from moving in between these two stable fixed points. It is important to note that in the case with intrinsic spine dynamics, while single spines are largely fluctuating and sporadically moving around the small and large mean spine-volume fixed points, a cell assembly is stably maintained by the ensemble property of intra-group spines: here quantified as the mean intra-group spine volume.

These results suggest that intrinsic spine dynamics can normalize the synaptic strength distribution to a stereotypical shape in the presence of ongoing Hebbian plasticity, and at the same time, enable the circuit to stably retain memory patterns in the form of cell assemblies with a bi-stable mean intra-cell assembly spine volume. The relative amplitudes of STDP and intrinsic spine dynamics are key parameters to achieving this bi-stability. For example, if STDP is too strong, relative to intrinsic spine dynamics, the small mean spine volume fixed point at 0.15 μm^3^ would disappear. We have already seen in our model that in the absence of intrinsic spine dynamics, the spine volume distribution sharply peaks at around the large mean spine volume fixed point and activity becomes too high (Fig. 1). Instead, if STDP is too weak, relative to intrinsic spine dynamics, the large mean spine volume fixed point around 0.55 μm^3^ could disappear. Below, we explore more generally what might go wrong with excessively strong intrinsic spine dynamics.

Interestingly, experimental results suggest that intrinsic spine dynamics are abnormally high in a mouse model of fragile X syndrome, *fmr1*KO (Nagaoka et al. 2016; Pan et al. 2010). Below, we constrain the parameters α and β of the diffusion coefficient *D*(*v*) = (α*v* + β)^2^/2 to characterize intrinsic spine dynamics in *fmr1*KO mice based on reported observations. Spine turnover is about twice as high in *fmr1*KO mice as in WT mice (Pan et al. 2010). Remarkably, this elevated spine turnover largely remains even when calcium activity is pharmacologically blocked, suggesting that this is due to abnormal intrinsic spine dynamics (Nagaoka et al. 2016). Another constraint is the spine volume distribution. Experimental reports comparing the spine volume distribution in *fmr1*KO and WT mice have mixed observations -- some studies detected more immature spines in *fmr1*KO but others detected no significant difference (He and Portera-Cailliau 2013). For simplicity, we assume that any differences in the spine volume distribution between *fmr1*KO and WT mice are negligible. Based on the numerical fitting to the observed spine turnover, these two constraints specify the parameters *a* = 0.43 day^−½^ and β = 0.021 μm^3^ day^−½^ for *fmr1*KO mice (Fig. 5; see also Fig. S3 for a systematic parameter search). Intuitively speaking, α and β are twice as high as the corresponding WT values. This is because, despite the presence of STDP, the spine volume distribution is largely set by the equilibrium distribution of the intrinsic spine dynamics, *P_eq_(v*) ∝ ı/(_v_ + β/α)^2^. It suggests that the ratio β/α should be similar between WT and *fmr1*KO to have similar spine volume distributions, and β in *fmr1*KO is twice as large as WT to account for the doubled spine turnover rate.

**Figure 5.**
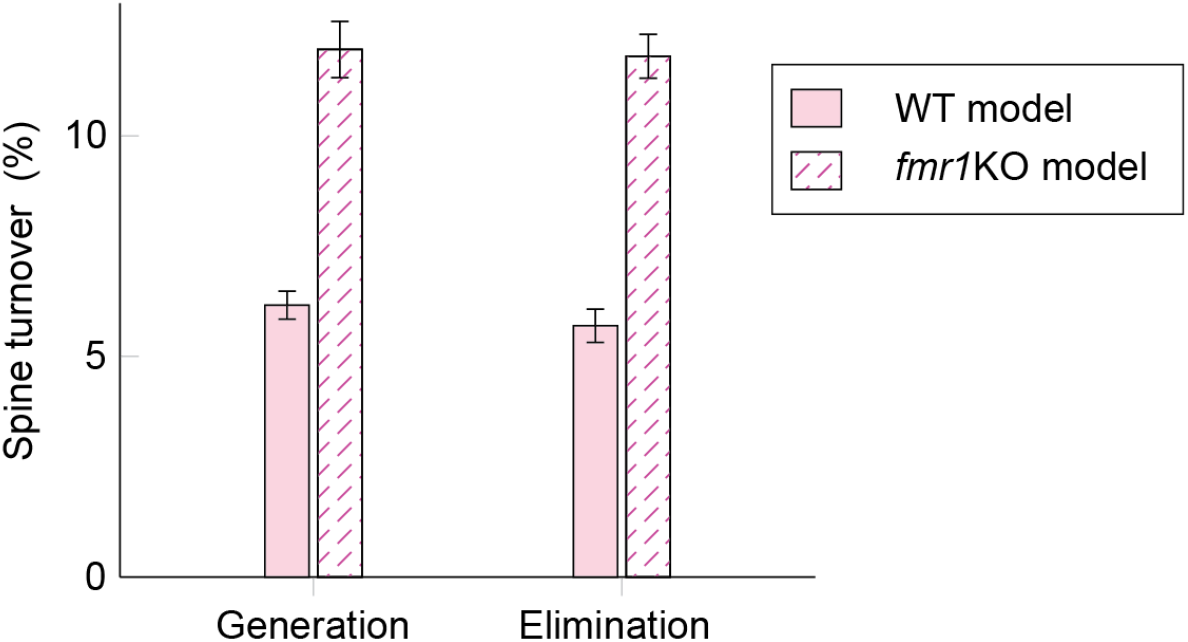
Spine turnover in our WT and *fmr1*KO models with two different levels of intrinsic spine dynamics.

Learning and memory deficits have been reported in *fmr1*KO mice (Padmashri et al. 2013). We investigated whether these learning and memory deficits could potentially be explained by the abnormal intrinsic spine dynamics modeled above. Consistent with the experimental observations, storing memory patterns was impaired in our *fmr1*KO model (Fig. 6A-H), where coherent spontaneous activation of cell assemblies rapidly faded after learning (Fig. 6B-D). This is because the mean spine volumes of the stimulated groups decreased during the maintenance period (Fig. 6F). This effect can be intuitively understood from the result in Fig. 4 -- memory cannot be stably stored in a cell assembly if intrinsic spine dynamics are too strong, relative to multiplicative STDP, because the stable fixed point of the mean intra-group spine volume around 0.55 μm^3^ disappears. Individual spines fluctuated as in the WT model but with a greater amount per unit time (Fig. 6G). As a result of excessively strong intrinsic spine dynamics and the decay of mean intra-group spine volume for the stimulated groups, the stimulated spines scattered around the spine volume distribution (Fig. 6E). This result is in contrast to the WT result, where stimulated spines are localized nearer the tail of the spine volume distribution (c.f. Fig. 3E). Similarly to the WT model, the external stimulation at the onset of learning increased the spine generation rate by about 2.5% without significantly altering the elimination rate. Note that the baseline turnover in this *fmr1*KO model was about twice as high as the WT model, consistent with the experimental observation (Nagaoka et al. 2016).

**Figure 6.**
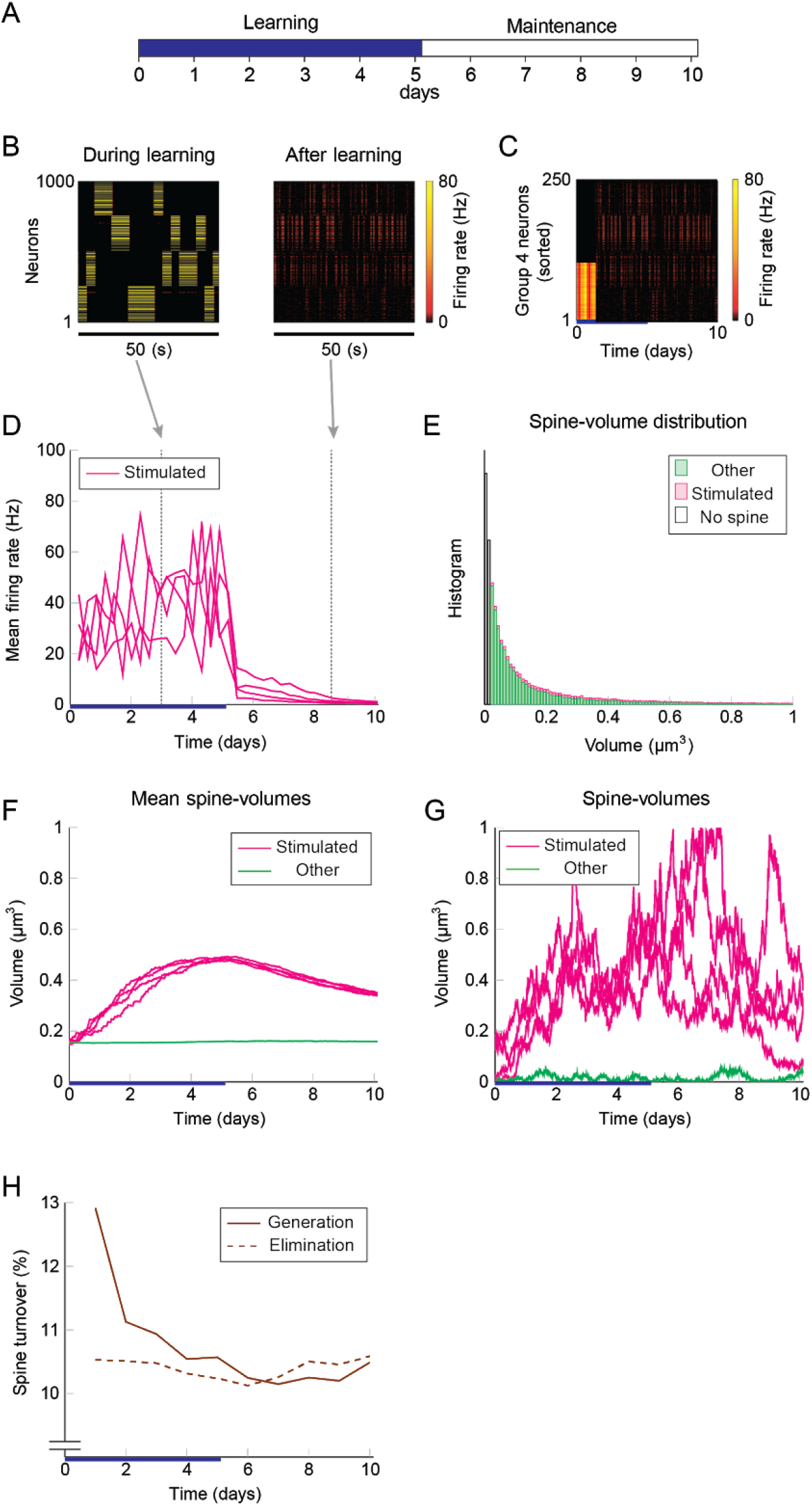
Network behavior in the presence of predicted *fmr1*KO intrinsic spine dynamics. (A-H) Conventions are as in Fig. 3.

The learning and memory deficits reproduced above are however potentially more severe than observed experimentally because *fmr1*KO mice have *some* learning capability, albeit limited. Therefore, we considered whether there may be some compensating mechanism, via which animals with excessively strong intrinsic spine dynamics rescue some learning and memory performance. We consider the regulation of potential connectivity as one candidate mechanism. Figure 7 explores how the stability of cell assemblies changes in our WT and *frm1*KO models (i.e. the simulations of Figs. 3 and 6) if the peak potential connectivity (Fig. 1B_1_) is systematically altered. A cell assembly in either the WT or *fmr1*KO model can exhibit one of the following three scenarios: its mean firing rate either (1) explodes (≥100 Hz) due to unstable learning, (2) fades (≤1 Hz), or (3) is stably maintained within a physiological range (>1 Hz and <100 Hz) during the maintenance period. The peak potential connectivity of 10.4% was already optimized for the WT model, such that nearly all cell assemblies are stable. The frequency of fading assemblies increased if potential connectivity was too low and that of exploding assemblies increased if potential connectivity was too high. However, the WT model stably maintained cell assemblies over a range of the peak potential connectivity from 9.4% to 10.8%. In contrast, the maintenance of cell assemblies in the *fmr1*KO model was much more sensitive to potential connectivity. As indicated in Fig. 7, most of the assemblies fade with the WT peak connectivity of 10.4%. The stability in the *fmr1*KO model first improved up to ~50% maintenance rate, but soon started to decline again due to exploding assemblies as potential connectivity increases. Given this result, it is an interesting possibility that *fmr1*KO mice with excessively strong intrinsic spine dynamics may locally up-regulate potential connectivity, relative to WT mice, to avoid catastrophic forgetting (Fig. 7 Right; marked as compensation). This hypothesis, which predicts an elevated frequency of exploding assemblies in *fmr1*KO mice, is consistent with the experimentally observed hyper neural activity and synchrony in *fmr1*KO mice (Gonçalves et al. 2013).

**Figure 7.**
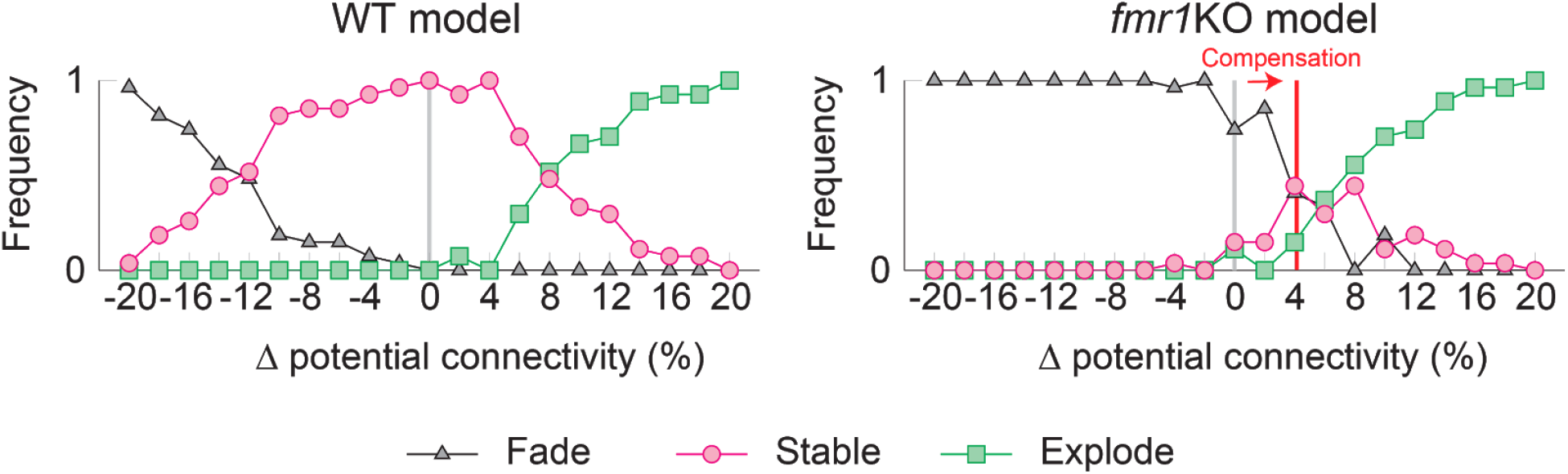
Varying the probability of recurrent excitatory to excitatory connections in the WT model (Left) and the *fmr1*KO model (Right) permits an increased chance of cell assembly fade or explosion at different connectivity levels. We hypothesize that *fmr1*KO mice may have a compensatory increase in potential connectivity (red line) to partially rescue stable cell assemblies.

## Discussion

In previous models of synaptic plasticity, changes in synaptic strengths are assumed to be activity-dependent (Poo et al. 2016; Mongillo et al. 2017). Recent experimental observations suggest that this is not the case. In this work, we modeled how activity-independent intrinsic spine dynamics (Yasumatsu et al. 2008; Nagaoka et al. 2016) affect learning and memory in recurrent circuit models. For simplicity, we assumed that spine volume is proportional to synaptic strength (Matsuzaki et al. 2004; Harvey and Svoboda 2007; Bosch et al. 2014). Contrary to a view that noisy synaptic changes are harmful to memory, intrinsic spine dynamics in our model play a positive role in preventing Hebbian-plasticity-driven non-specific growth of synapses. Specifically, as a result of the interaction between STDP and intrinsic spine dynamics in our model, the mean volume of a cell assembly’s spines exhibits bistability, which is suitable to sustain memory. STDP keeps spines within a cell assembly due to the coherent spontaneous activity of the membership neurons (e.g. (Wei and Koulakov 2014; Diekelmann and Born 2010; Kenet et al. 2003), and intrinsic spine dynamics constitutively normalize all spines toward a physiological distribution.

In our model, memory is maintained by the total strength of synapses that connect neurons within an assembly. Individual synapses fluctuate and exhibit constant turnover, but this does not degrade the memory as long as the net strength is kept. However, one property of the current model is that, given the innate Hebbian instability of cell assemblies in spontaneously active recurrent networks, highly overlapping cell assemblies likely merge during maintenance, even in the presence of intrinsic spine dynamics. In this sense, either a preprocessing mechanism that decorrelates memory patterns (Perez-Orive et al. 2002; Leutgeb et al. 2007) or a more elaborate additional mechanism that stabilizes individual synapses (Frey and Morris 1997; Ziegler et al. 2015; Benna and Fusi 2016) is helpful for improving the memory capacity (Hopfield 1982).

There are several experimental observations that support the role of intrinsic spine dynamics in shaping the spine volume distribution. First, the spine volume distribution in the absence of neural activity and plasticity was well predicted by intrinsic spine dynamics (Yasumatsu et al. 2008), and the distribution is similar in the presence of neural activity. Second, the spine volume distribution was robust to experimental manipulations to synaptic plasticity. Remarkably, calcineurin KO mice with little LTD (Okazaki et al. 2018) exhibited a spine volume distribution that was similar to WT mice. This raises an argument against the hypothesis that the spine volume distribution is set by the balance between LTP and LTD. We suggest that previous computational models that do not include intrinsic spine dynamics miss an important component underlying synaptic organization.

We studied the interplay between activity-dependent synaptic plasticity and intrinsic spine dynamics in the formation and maintenance of cell assemblies in recurrent networks. Conventional studies of intrinsic spine dynamics often focus on independent synapses (Yasumatsu et al. 2008) or learning in a local feedforward network (Matsubara and Uehara 2016) instead of learning in a recurrently connected network. Another study explored the consequence of volatile synaptic strengths in recurrently connected networks without studying how such volatility affects activity-dependent plasticity (Mongillo et al. 2018). Other studies model stochastic changes of synaptic strength (Loewenstein et al. 2011) or connectivity (Deger et al. 2012; Fauth et al. 2015) without distinguishing activity-dependent and -independent parts. Hence, these works do not distinguish their separate roles in memory and synaptic normalization. Another line of studies (Zenke et al. 2015; Litwin-Kumar and Doiron 2014) model activity-dependent synaptic plasticity to account for the formation and maintenance of cell assemblies in a recurrent network. However, because these models do not include intrinsic spine dynamics, their synaptic strength distributions are typically sensitive to the fine balance between LTP and LTD and counter to the experimental observations described above. Thus, the proposed model postulates how synaptic normalization, by intrinsic spine dynamics, maintains most synapses that are not participating in a cell assembly weak. In addition, intrinsic spine dynamics in our model work as a homeostatic mechanism that stabilizes Hebbian plasticity, although they are activity independent and an explicit sensing-and-control (Shah and Crair 2008; Davis 2006) mechanism is absent. For this to work, the relative magnitude of Hebbian plasticity and intrinsic spine dynamics is important; For example, this stabilization fails if Hebbian plasticity is too fast (see Fig. 4). While we conjecture that intrinsic spine dynamics can stabilize slow Hebbian plasticity in adults, other fast forms of homeostatic plasticity, such as inhibitory plasticity (Vogels et al. 2011; Litwin-Kumar and Doiron 2014), might be helpful in the young, or once neural activity deviates beyond the level that intrinsic spine dynamics can compensate for.

Another main contribution of this study is the relation between intrinsic spine dynamics and ASD. The observation that baseline spine turnover is abnormally high in various ASD mouse models (Isshiki et al. 2014), including a model animal for fragile X syndrome (*fmr1*KO) (Pan et al. 2010) suggests abnormal intrinsic spine dynamics could be one candidate substrate of ASD. Indeed, a recent experiment has confirmed that high baseline spine turnover in *fmr1*KO mice is activity-independent (Nagaoka et al. 2016), and intrinsic spine dynamics are stronger in *fmr1*KO (Ishii et al. 2018). Based on these experimental observations, we fitted parameters characterizing intrinsic spine dynamics in *fmr1*KO mice and found that they explain the learning deficit observed in *fmr1*KO mice (Padmashri et al. 2013). Interestingly, when we included a compensatory increase in recurrent excitatory connectivity to rescue some memory, the model reproduced epileptic-like neural activity that has been reported experimentally in *fmr1*KO mice (Musumeci et al. 2000). More generally, the disrupted cortical connectivity theory of ASD (Courchesne and Pierce 2005; Kana et al. 2011) argues that deficiency of cortical long-range connections and compensatory strengthening of local connections is a general feature of ASD. We contend that because there are typically fewer long-range connections, which therefore limits the positive feedback effect of Hebbian plasticity that is required for maintaining cell assemblies, they are especially susceptible to degradation due to excessively strong intrinsic spine dynamics, such as in our *fmr1*KO model. Furthermore, the learning deficiency in our proposed local cortical circuit model of *fmr1*KO, and its rescue by a compensatory increase in the local connectivity (Fig. 7), is consistent with this theory. Hence, it is an intriguing possibility that pharmacological manipulations (Nagaoka et al. 2016), or a future neurofeedback technology (Ganguly and Poo 2013; Yahata et al. 2016), could be used to rescue memory and learning, and long-range neural association, by reducing intrinsic spine dynamics or producing network motifs (Watanabe and Rees 2015) efficiently connecting target brain areas.

A similar concept to intrinsic spine dynamics that has been used to describe ASD is intrinsic forgetting (Davis and Zhong 2017). These two mechanisms are both related to chronic molecular signaling, which slowly degrades synapses and therefore memories. However, important differences between the two are how they are regulated in ASD animals and how they could possibly be involved in producing flexible behavior. A decrease in intrinsic forgetting has been argued to disable flexible behavior by maintaining conflicting memories (Davis and Zhong 2017; Reaume et al. 2011). In contrast, the excessively strong intrinsic spine dynamics found in ASD animals (Isshiki et al. 2014; Nagaoka et al. 2016; Pan et al. 2010) would work in the opposite manner, by facilitating forgetting. Namely, stable cell assemblies in our *fmr1*KO model, typically require more neurons than the WT model to counter excess synaptic normalization. Hence, given a hypothesis that the number of total neurons is roughly the same between *fmr1*KO and WT mice, *fmr1*KO mice may afford a smaller number of cell assemblies in the brain, possibly reducing the number of behavioral repertoires.

Overall, our spine dynamics model provides a novel path relating spine statistics, memory, and ASD. Notably, it is currently difficult to block intrinsic spine dynamics experimentally *in vivo* without affecting plasticity, because they are thought to be caused by the thermal agitation of molecules. This underscores the importance of a modeling study. Selective manipulations of intrinsic spine dynamics are an intriguing candidate direction to influence memory and learning, and the current model serves as a guide.

## Methods

### Network and neurons

A local network of *N_E_* excitatory *N_I_* and inhibitory leaky-integrate-and-fire neurons (Fig. 1A; see e.g., (Tuckwell 1988)) is considered. We simulate a network with (*N_E_*, *N_I_*) = (1000, 200). The dynamics of membrane potential *V_i_* of neuron *i* is described by

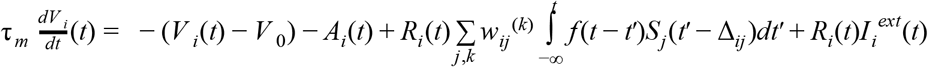

with membrane time constant τ*_m_* = 20 ms, resting potential *V*_0_ = − 70 mV, adaptation *A_i_*, refractory coefficient *R_i_*, *k* th (*k* = 1, 2, …), synaptic strength *w_ij_*^(*k*)^ from neuron *j* to neuron *j*, spike train 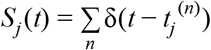 of neuron *j* composed of *n* th (*n* = 1, 2, …) spike time *t_j_*^(*n*)^, random axonal delay Δ*_ij_* from neuron *j* to neuron *i* uniformly sampled from interval [0.5, 5.0] ms, and external input 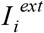 to neuron *i* (see below). Note that δ is the Dirac delta function. The time course of postsynaptic input is modeled using the alpha function (Gerstner et al. 2014),

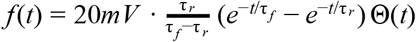

with rise time τ*_r_* = 0.5 ms, fall time τ*_f_* = 2.0 ms, and the Heaviside step function Θ(0 that takes 1 for *t* ≥ 0 and 0 for *t* < 0. The peak value of *f*(*t*) is about 0.39 mV. A spike is emitted when *V_i_* reaches a spiking threshold at − 50 mV and then *V_i_* is reset to *V*_0_. Excitatory neurons receive adaptation input *A_i_* that reflects a slow Na^+^-activated K^+^ current (Wang et al. 2003) (Fig. 1B_4_), with dynamics described by

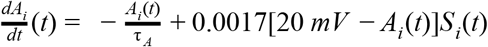

with time constant τ*_A_* = 13 s. In contrast, we assume no adaptation (*A_i_* = 0) in inhibitory neurons. Refractoriness is imposed by *R_i_* . *R_i_* is fixed at 0 after each spike of neuron *i* for 1 ms (absolute refractoriness), and then recovers toward 1 following

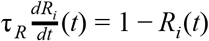

with time constant τ*_R_* = 3.5 ms (relative refractoriness). The differential equations are numerically solved using the Euler method with bin size Δ*t* = 0.1 ms.

### Network topology

In the network model of Fig. 1A, excitatory neurons, *E*, and inhibitory neurons, *I*, are sparsely connected. The connections from excitatory to inhibitory neurons (*E* → *I*) and those from inhibitory to excitatory neurons (*I* → *E*) are randomly generated with 10% connection probability. Each of these directed connections is mediated by a single synapse, whose strength is randomly drawn from a uniform distribution in the range [0, 31] mV for *E* → *I* and [-31, 0] mV for *I* → *E*, respectively. According to our neuron model described above, a typical *E* → *I* or *I* → *E* synapse produces a postsynaptic potential of 6 mV. For simplicity, we assume no direct connections between inhibitory neurons. Excitatory neurons are equidistantly placed on a one-dimensional ring of circumference 1 a.u. that represents the feature space. Note that the tuning distance in feature space *d_fs_* does not necessarily correspond to the physical location of a neuron, or the distance between any two neurons. The potential connections from excitatory to excitatory neurons (*E* → *E*) are randomly generated according to the Gaussian probability profile (10.4% * exp[−0.5 * (*d_fs_*/0.1)^2^], Fig. 1B_1_), which peaks at 10.4% for similarly tuned neurons (*d_fs_* ≈ 0), and decays toward 0 with a length-constant of 0.1 as *d_fs_* increases. Although this *E* → *E* peak connection probability is optimized for memory retention, simulation outcomes are robust with this parameter in our model (see, Fig. 7, WT model). Effectively, this allows excitatory neurons closer in the feature space to be more interconnected, and those farther apart to be less so. If there is a potential connection from one neuron to another, we randomly assign a fixed number *K* (*K* = 1,2, …, 10) of spines from a Poisson distribution

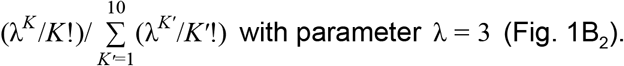

### Stimulation

In addition to the recurrent input, neuron *i* also receives spike train 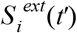 as external input (dashed connection in Fig. 1A). At baseline, 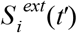 is generated by a Poisson process with firing intensity 60 Hz. Input neurons are not modeled explicitly here. The external input to each neuron is given by 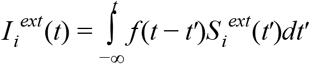. This sets the baseline membrane potential and firing rate of neurons to −58.6 ± 2.4 mV and 0.13 ± 0.08 Hz, respectively.

We divide the feature space of excitatory neurons into 4 equally-sized consecutive parts and randomly select 40% of the neurons from each part as a stimulated neuron group (Fig. 1A). During the learning period (indicated by blue horizontal bars below the time axes in Figs. 2, 3, and 6), one of the 4 stimulated neuron groups is randomly selected with probability ¼ and receives additional Poisson spikes at 750 Hz for 3 s. During the learning period, all inhibitory neurons also receive additional Poisson spikes at 300 Hz. The learning period starts at *t* = 0 and ends when at least one group’s mean intra-group spine volume reaches ≥ 0.49 μm^3^.

### Spine dynamics

Unlike *E* → *I* and *I* → *E* synapses, which have fixed weights, *E → E* synapses (orange and brown dashed in Fig. 1A) change in time via two independent mechanisms: STDP and intrinsic spine dynamics. We assume that synaptic strengths for *E → E* synapses are essentially proportional to their spine volumes and therefore model their spine volume dynamics as changes in synaptic strengths. Changes in the *k* th (*k* = 1, 2, …, *K_ij_*) spine volume *v_ij_*^(*k*)^ on neuron *i*, receiving signal from neuron *j*, is modeled by

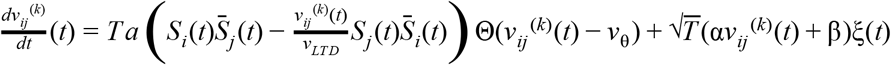

where the first and second terms on the right hand side describe changes by multiplicative STDP (van Rossum et al. 2000) and intrinsic spine dynamics (Yasumatsu et al. 2008) respectively. *T* is a speed-up factor that we describe below, *a* = 7.6 · 10 ^9^ μm^3^ is the amplitude of STDP, 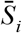 is the running average of past spiking activity of neuron *i*, i.e.,

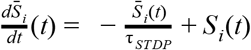

with averaging time constant τ*_STDP_* = 20 ms, and *v_LTD_* = 0.5 μrrr is the scaling factor for volume-dependent LTD (van Rossum et al. 2000). STDP is assumed to be absent for small spines of *v_ij_*^(*k*)^ < *v*_θ_ = 0.02 μm^3^. Slope parameter α = 0.2 day^−½^ and offset parameter β = 0.01 μm^3^ day^−½^ for intrinsic spine dynamics are set as previously experimentally observed (Yasumatsu et al. 2008). ξ is white noise with the autocorrelation function < ξ(*t*)ξ(*t′*) >= δ(*t-t′*). The above Langevin equation is numerically solved by the Euler method with bin size Δ*t* = 0.1 ms. In addition, we set reflecting boundaries for spine volume to enforce 0 ≤ *v_ij_*^(*k*)^ ≤ 1.0 μm^3^ for all spines. One problem is that it is too time consuming to directly simulate the 10-day learning period studied with the fine time resolution required to simulate STDP and intrinsic spine dynamics. We therefore run *T* = 3.3 · 10^4^ times shorter simulations by speeding up both STDP and intrinsic spine dynamics by factors *T* and 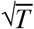, respectively. (Note that volume changes *v*(*t* + Δ*t*) − *v*(*t*) by intrinsic spine dynamics are diffusive and scale with the square root of time duration 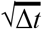.) This way, we can extrapolate spine volume changes happening during 10 days based on shorter simulations up to 3000 s. We display the time before this conversion in panels describing neural activity in seconds, but display the time after this conversion in panels describing learning in days. We initially set *E → E* spine volumes by randomly sampling from the equilibrium distribution *P_ss_*(*v*) ∝ (α*v* + β)^−2^, which is set by the intrinsic spine dynamics. Synaptic strength *w* of a spine with volume *v* is then assumed to be

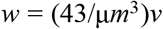

for *v* ≥ *v*_θ_ (a functional spine) and 0 for *v* < *v*_θ_ (a non-functional spine, e.g., filopodia). The median spine volume of *P_ss_*(*v*) is 0.047 μm^3^ and such a spine produces 0.8 mV of excitatory postsynaptic potential. We set v_θ_ to be a threshold volume typically used in experiments to detect spines (Yasumatsu et al. 2008). We define spine gain and loss by a fraction of spines passing this threshold from below and above per day, respectively. The exact value of *v*_θ_ does not matter for our results as long as it is sufficiently small.

### Itô vs Stratonovich interpretation

The meaning of *fluctuation* is different under the Itô and Stratonovich interpretations of intrinsic spine dynamics (Gardiner 1985). The intrinsic spine dynamics under the Itô interpretation that we study in the main text are described by

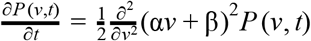

In this view, changes in spine volume distribution are purely produced by the volume-dependent *fluctuation* with amplitude α*v* + β. In contrast, the Stratonovich interpretation represents the same equation in a different way:

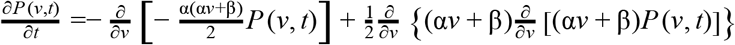

In this view, changes in spine volume distribution are described by two terms: the first term is the drift term produced by apparent force α(α*v* + β)/2 and the second term is produced by the volume-dependent *fluctuation αv +* β. Hence, while the above two equations are identical, there is a semantic difference regarding what *fluctuation* means. According to the Itô interpretation, only the *fluctuation* drives spine volume changes and this *fluctuation* preserves mean spine volume (*i.e*., martingale (Øksendal 2000)) except for a boundary effect. According to the Stratonovich interpretation, the apparent force shrinks and the *fluctuation* enlarges mean spine volume respectively, and the two effects are cancelled. These two interpretations become the same in the special case of α = 0, namely, when the *fluctuation* is volume independent.

### Simulation environment

Simulations were performed in custom written C code with the forward Euler-integration method and a step size of 0.1 ms. Post-simulation analysis was undertaken with MATLAB. Source code is available upon request.

## Author contributions

HK and TT conceived the project. JH and KH performed numerical simulations. JH, KH, HK, and TT analyzed the results. JH, KH, HK, and TT drafted and revised the manuscript.

## Supplementary material

**Figure S1.**
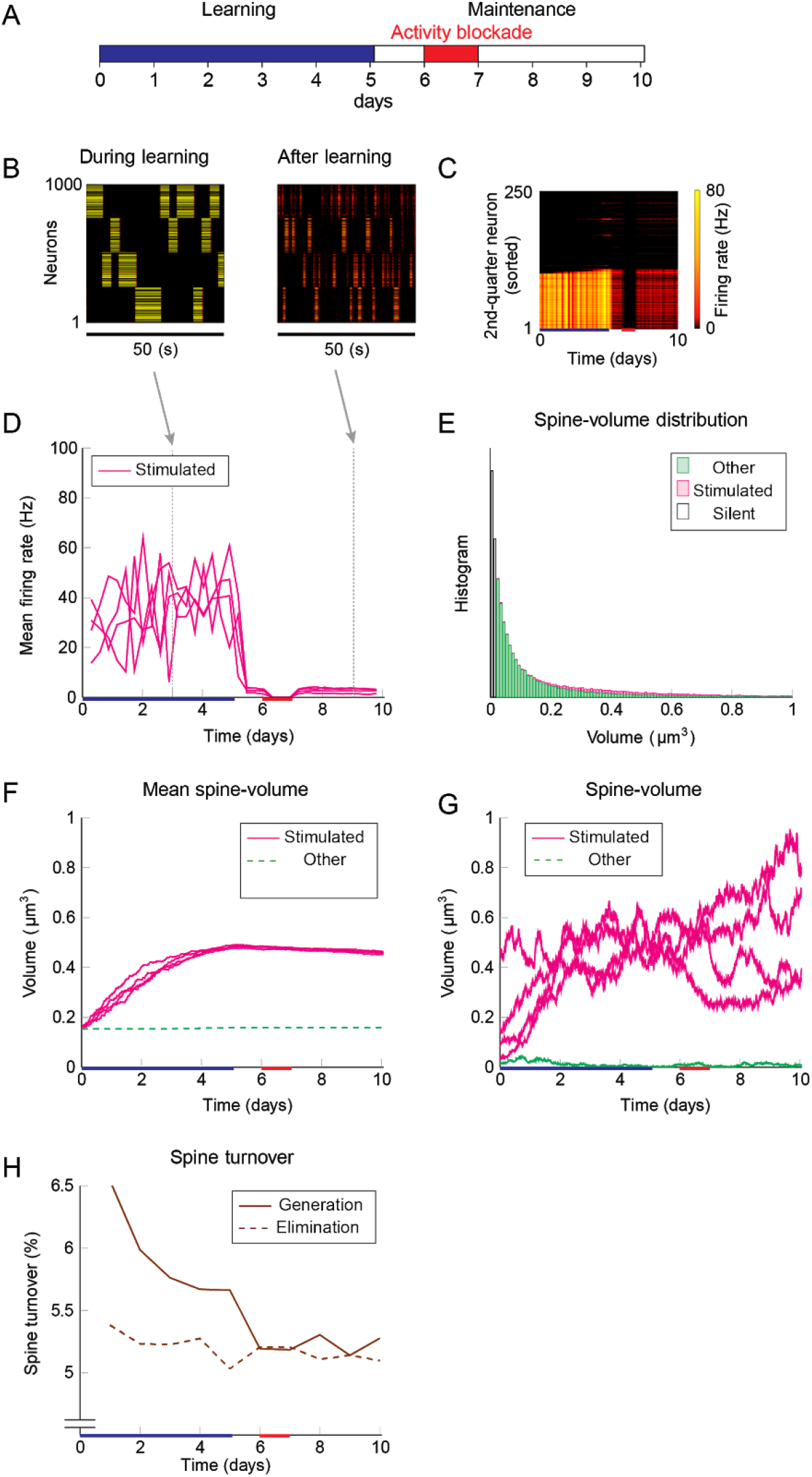
Network behavior in the presence of intrinsic spine dynamics, similar to Fig 3, but with 1 day blockade of neural activity during the maintenance period. The activity blockade does not change the results. (A-H) Conventions are as in Fig. 3.

**Figure S2.**
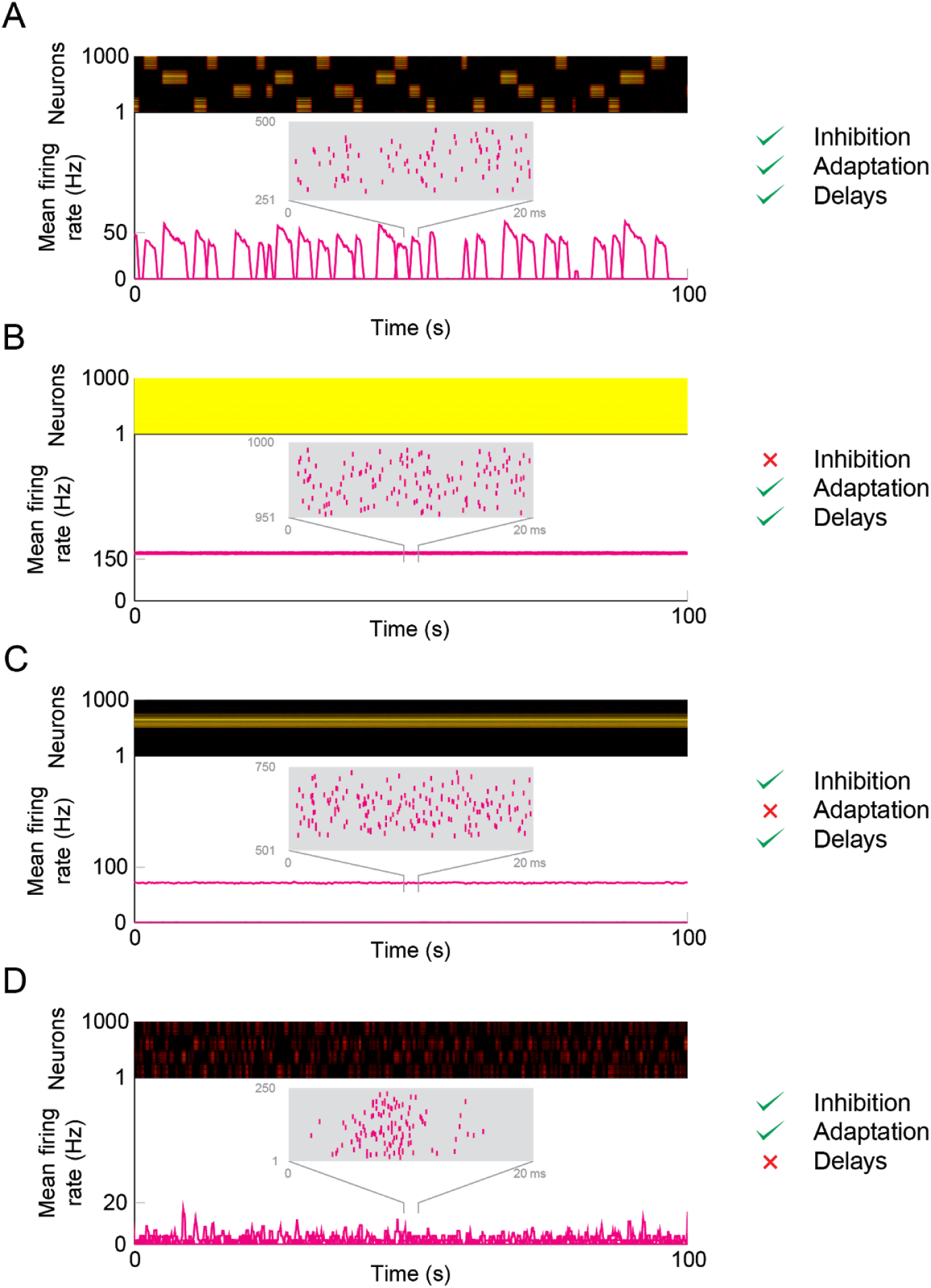
Description of different mechanisms in the model. (A) With inhibition, adaptation, and axonal delays all functioning, the network retains all cell assemblies where each assembly is rehearsed for several seconds during spontaneous activity. (B) When the inhibitory neurons are removed all excitatory neurons continuously fire a saturated rate >150 Hz. (C) When adaptation is removed from excitatory neurons, only one assembly dominates. (D) When axonal delays are removed, the four memories are somewhat maintained, albeit with very fast noisy switching and a much lower firing rate. The removal of a mechanism was done after learning and at the onset of the maintenance period, with all other mechanisms, including STDP and intrinsic spine dynamics, functionally preserved.

**Figure S3.**
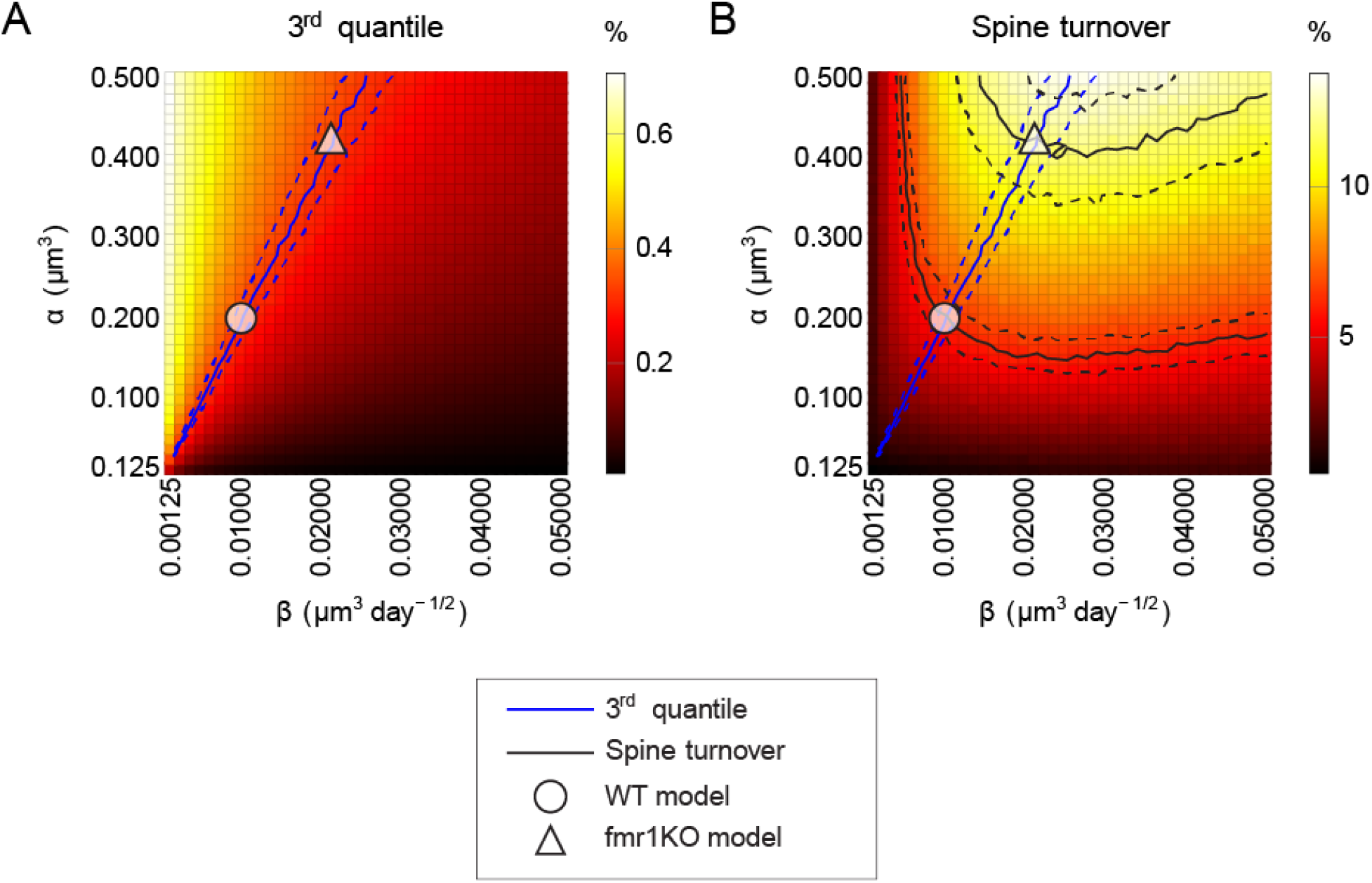
Systematic exploration of intrinsic spine dynamics’ parameters α and β. (A) The 3rd quantile of the equilibrium spine volume distribution is shown in color as a function of α and β. The entire distribution roughly scales with the ratio α/β as expected based on the theoretical consideration. The blue solid line (and dashed lines) indicates the experimentally observed 3rd quantile (and ±10% range). (B) Spine turnover is shown in color as a function of parameters α and β. Increases in either α or β result in increases in spine turnover. The two solid black lines (and dashed lines) indicate experimentally observed spine turnover for WT and *fmr1*KO animals (and ±10% range). We therefore used the two parameter combinations of α and β at the cross points of the black and blue solid lines in our WT and *fmr1*KO models.

